# Temporal modulation of the NF-κB RelA network in response to different types of DNA damage

**DOI:** 10.1101/2020.08.11.246504

**Authors:** Amy Campbell, Catarina Ferraz Franco, Ling-I Su, Simon Perkins, Andrew R. Jones, Philip J. Brownridge, Neil D. Perkins, Claire E. Eyers

**Author notes:** These authors contributed equally to this manuscript (Amy Campbell, Catarina Ferraz Franco).

## Abstract

Different types of DNA damage can initiate phosphorylation-mediated signalling cascades that result in stimulus specific pro- or anti-apoptotic cellular responses. Amongst its many roles, the NF-κB transcription factor RelA is central to these DNA damage response pathways. However, we still lack understanding of the co-ordinated signalling mechanisms that permit different DNA damaging agents to induce distinct cellular outcomes through RelA. Here, we use label-free quantitative phosphoproteomics to examine the temporal effects of exposure of U2OS cells to either etoposide (ETO) or hydroxyurea (HU) by monitoring the phosphorylation status of RelA and its protein binding partners. Although few stimulus-specific differences were identified in the constituents of phosphorylated RelA interactome after exposure to these DNA damaging agents, we observed subtle, but significant, changes in their phosphorylation states, as a function of both type and duration of treatment. The DNA double strand break (DSB)-inducing ETO invoked more rapid, sustained responses than HU, with regulated targets primarily involved in transcription, cell division and canonical DSB repair. Kinase substrate prediction of ETO-regulated phosphosites suggest abrogation of CDK1 and ERK1 signalling, in addition to the known induction of ATM/ATR. In contrast, HU-induced replicative stress mediated temporally dynamic regulation, with phosphorylated RelA binding partners having roles in rRNA/mRNA processing and translational initiation, many of which contained a 14-3-3ε binding motif, and were putative substrates of the dual specificity kinase CLK1. Our data thus point to differential regulation of key cellular processes and the involvement of distinct signalling pathways in modulating DNA damage-specific functions of RelA.

## INTRODUCTION

RelA, also known as p65, is a key member of the NF-κB family of transcription factors, which serve as ‘master regulators’ of the cellular inflammatory and stress responses, and are key components that maintain tissue homeostasis and contribute to aging. As part of a functional homo- or hetero-dimer (preferentially with p50), RelA regulates a diverse set of genes involved in core cellular processes such as inflammation, proliferation, apoptosis and metastasis [1]. The precise functional roles (the complement of genes transcribed) of RelA are governed both by its post-translational modification (PTM) status, and the transcriptional complexes formed, which are themselves regulated in a co-ordinated fashion through numerous cell signalling pathways [2–4]. PTMs have the potential to regulate many aspects of NF-κB signalling in response to different stimuli, including nuclear translocation, protein-protein or protein-DNA interactions, and stability, often working in a combinatorial fashion to regulate complex formation and transcriptional output.

While NF-κB signalling is commonly associated with inflammation and the immune response, these pathways also play key roles in the cellular response to DNA stress. Consequently, aberrant NF-κB signalling is observed in, and often a contributing factor to, many human diseases, including autoimmune disorders, chronic inflammatory diseases, and cancer [1,5–7]. NF-κB signalling is also central to age-related degenerative diseases as a result of accumulated age-related DNA damage [8]. The pro-survival effects mediated by NF-κB in response to specific-types of DNA damage in part explains the cellular resistance to chemotherapy. Hence, a clear understanding of NF-κB signalling in response to DNA damage is important, not only in the context of ageing, but also to enhance cancer treatment strategies.

In response to different types of DNA damage, cells invoke specific responses that promote DNA repair, induce cell cycle arrest, regulate cell death or induce cellular senescence [9,10]. For example, inhibition of DNA replication using hydroxyurea (HU), which inhibits ribonuclease diphosphate reductase, arrests cells primarily in S-phase [11–15]. In contrast, agents such as etoposide (ETO), which induces DNA double stranded breaks (DSBs) by preventing topoisomerase II-mediated DNA re-ligation during replication and cell division, cause cells to accumulate in the G2/M phase of the cell cycle [16–19]. Whilst the activation of NF-κB-responsive genes following DNA stress have been investigated, the mechanisms of regulating these differential outputs remain poorly characterised [1,5–7]. Both HU and ETO invoke a DNA damage stress response through ataxia telangiectasia mutated (ATM)-mediated activation of NF-κB, leading to elevated levels of NF-κB targets such as Survivin and Bcl-xL. However, these two types of DNA damaging agents result in distinct cellular responses in a variety of tumor-derived cell lines and in primary cells cancer cells, in part due to induction of different NF-κB gene expression patterns [19–22]; HU mediates a pro-apoptotic response, while an anti-apoptotic response results following cellular exposure to ETO [22]. Differential involvement of p53 in regulating NF-κB outputs in response to these different types of DNA damaging agents also plays a role: while the transcriptional response to HU requires p53 for both early and late NF-κB target gene expression, NF-κB-mediated transcription in response to ETO is independent of p53 [22].

The activity of all NF-κB subunits, including RelA, is extensively controlled by a variety of PTMs including: phosphorylation, acetylation, methylation, glycosylation, proline isomerisation, cysteine oxidation and ubiquitination [23–26]. Of these, phosphorylation accounts for the majority of the well-understood dynamic mechanisms of regulation, both direct *e.g*. pSer45 modulation of DNA binding [26], and indirect *e.g*. pSer276, which controls a number of other PTMs, including ubquitination and thus RelA stability [2,27,28]. Of relevance here, Thr505 phosphorylation has been shown to mediate pro-apoptotic effects upon cisplatin-induced DNA damage [29,30]. Given the complexity of NF-κB signalling, unravelling the mechanisms by which NF-κB subunits, particularly RelA, induce differential transcriptional specificity in a context-dependent manner is challenging.

The sensitivity and versatility of mass spectrometry (MS) makes it ideal for the study (identification and quantification) of dynamic PTMs and the regulated binding partners of transcription factors and their transcriptional complexes. Here we utilise a label free phosphoproteomics approach to map DNA damage-induced changes in RelA binding partners and their phosphorylation status, in response to either HU-induced replicative stress, or ETO-mediated DSB. We find that whilst RelA associated proteins remain largely unchanged as a function of DNA stress, there are notable changes in the dynamics of phosphorylation of the RelA-interactome as a function of treatment. These findings point towards different signalling pathways directing the formation of specific RelA networks dependent on the type of DNA damage that likely help direct transcriptional output and thus the cellular response.

## MATERIALS and METHODS

### Generation and culture of HA-RelA U2OS cells

The HA-RelA U2OS cell line was generated by transfecting wild-type U2OS cells with the plasmid pRcRSV HA RelA plasmid, which also expresses the neomycin gene. Cells stably expressing HA tagged RelA were then selected by treatment with G418 (600 µg/mL, Melford Chemicals, cat. no. G0175), with pooled populations of cells used in experiments.

### Cell treatment and lysis

U2OS cells expressing HA-RelA were cultured in DMEM supplemented with 10% (v/v) fetal bovine serum, penicillin (100 U/mL), streptomycin (100 U/mL), and L-glutamine (2 mM) at 37 °C, 5% CO2. Once 80% confluence was reached, cells were treated with either 50 μM etoposide (50 mM stock solution in DMSO) or 2 mM hydroxyurea (100 mM stock solution in DMEM, 5% DMSO), by addition to the culture media, for the indicated times. Control cells were grown for the maximum time (2 h) in 0.1% (v/v) DMSO. After removal of the media, cells were washed three times with ice cold PBS and collected in 400 μL of lysis buffer (100 mM Tris-HCl pH 8.0, 150 mM NaCl, 0.5 mM EDTA, 0.5% (v/v) Triton X-100, 0.5% (v/v) NP-40, 1 x PhosSTOP™ phosphatase inhibitor cocktail tablet (Roche), 1 x cOmplete™ Protease Inhibitor Cocktail (Roche), 50 U/mL of benzonase) into 1.5 mL microtubes using a cell scraper. Cells were then incubated on ice for 120 min to permit cell lysis to occur. Cell lysates were then cleared by centrifugation (16,000 x *g*, 10 min, 4°C) and protein concentration was determined using the Bradford assay. A total protein amount of 4 mg was set aside for HA-RelA immunoprecipitation.

### Immunoprecipitation of HA-RelA using Anti-HA magnetic beads

Pierce™ Anti-HA Magnetic Beads were washed with 0.05% (v/v) TBST and then with lysis buffer prior to addition of 4 mg cleared cell lysate (at a ratio of beads to protein of 1:2000). Beads and lysate were incubated overnight using an end-over-end rotor at 4°C. Beads were then collected using a magnetic stand and washed three times with 40 mM Tris-HCl pH 8.0, 0.1% (v/v) NP-40, 1 mM EGTA, 6 mM EDTA, 6 mM DTT, 0.5 M NaCl, 1 x PhosSTOP™ phosphatase inhibitor cocktail tablet (Roche), 1 x cOmplete™ Protease Inhibitor Cocktail (Roche); one time with HPLC grade water and finally, three times with 25 mM ammonium bicarbonate (AMBIC). To recover the immunoprecipitated material, the beads were resuspended in 100 μL of 25 mM AMBIC to which 6 μL of a 1% (w/v) solution of RapiGest (Waters, UK) in 25 mM AMBIC was added. Samples were then heated to 80 °C for 10 min. Supernatants containing the eluted proteins were recovered using a magnetic stand and used for in-solution digestion. For western blots the following antibodies were used: Anti-RelA (SC-372, Santa Cruz), Anti-HA (Merck, HA-7), Anti phospho-histone H2AX (2577, Cell Signalling).

### In-solution digestion and TiO_2_-based enrichment of phosphorylated peptides

Samples were digested with trypsin and subjected to TiO_2_-based phosphopeptide enrichment as previously described [31]. In brief, disulfide bonds were reduced by addition of DTT to a final concentration of 4 mM (10 min, 60 °C with gentle agitation). Samples were cooled to room temperature and free cysteines alkylated with 14 mM iodoacetamide (30 min, RT), prior to addition of DTT (to 7 mM final) to quench excess iodoacetamide. Samples were digested with trypsin (1:50 trypsin:protein ratio) overnight at 37°C with light agitation (450 rpm on a Eppendorf thermomixer). Digestion was stopped by the addition of trifluoracetic acid (TFA) to a final concentration of 0.5% (v/v) and incubated for 45 min at 37°C. RapiGest insoluble hydrolysis products were removed by centrifugation (13 000 x g, 15 min, 4°C). Digested samples were dried by vacuum centrifugation and resolubilized in TiO_2_ loading buffer (80% acetonitrile, 5% TFA, 1 M glycolic acid) to achieve a peptide concentration of 1 μg/μL and sonicated for 10 min. TiO_2_ beads (50 μg/μL in loading buffer) were mixed at 1400 rpm (with intermittent vortexing to prevent the beads from pelleting) with the resuspended peptides (bead:protein ratio of 5:1) for 20 min at RT. TiO_2_ beads were recovered by centrifugation (2000 x g, 1 min) and washed successively for 10 min (1400 rpm) with 150 μL of loading buffer, 80% acetonitrile 1% TFA, then 10% acetonitrile, 0.2% TFA. Prior to elution, beads with bound phosphopeptides were dried to completion by vacuum centrifugation. Bound phosphopeptides were then eluted by addition of 100 μL of 1% (v/v) ammonium hydroxide and then with 100 μL of 5% (v/v) ammonium hydroxide. Both elutions were combined and dried by vacuum centrifugation.

### Liquid chromatography-tandem mass spectrometry (LC-MS/MS) analysis

Phosphopeptide enriched samples were analysed by LC-MS/MS using an Ultimate 3000 RSLC™ nano system (Thermo Scientific, Hemel Hempstead) coupled to a QExactive HF mass spectrometer (Thermo Scientific). The samples were loaded onto a trapping column (Thermo Scientific, PepMap100, C18, 300 μm X 5 mm), using partial loop injection, for seven minutes at a flow rate of 4 μL/min with 0.1% (v/v) FA, and resolved on an analytical column (Easy-Spray C18 75 µm x 500 mm 2 µm column) using a gradient of 97% A (0.1% formic acid), 3% B (99.9% ACN 0.1% formic acid) to 60% A, 40% B over 80 minutes at a flow rate of 300 nL/min. Data-dependent acquisition employed a 60,000 resolution full-scan MS scan (MS1) with AGC set to 3e6 ions with a maximum fill time of 100 ms. The 10 most abundant peaks were selected for MS/MS using a 60,000 resolution scan (AGC set to 1e5 ions with a maximum fill time of 100 ms) with an ion selection window of 1.2 m/z and a normalised collision energy of 29. A 20 sec dynamic exclusion window was used to minimise repeated selection of peptides for MS/MS.

### LC-MS/MS data analysis

LC-MS/MS files were aligned in Progenesis QI for Proteomics label-free analysis software. At this stage, no normalisation was performed and an aggregate file containing raw abundances from all the peaks across all runs was exported from Progenesis. These files were searched against UniProt Human reviewed protein database (downloaded December 2015; 20,187 sequences) using MASCOT (v 2.6) within Proteome Discoverer (v. 1.4; Thermo Scientific). Parameters were set as follows: MS1 tolerance of 10 ppm, MS2 mass tolerance of 0.01 Da; enzyme specificity was defined as trypsin with two missed cleavages allowed; Carbamidomethyl Cys was set as a fixed modification; Met oxidation, and Ser/Thr/Tyr/His phosphorylation were defined as variable modifications. Data were filtered to a 1% false discovery rate (FDR) on peptide spectrum matches (PSMs) using automatic decoy searching with MASCOT. *ptm*RS node with Proteome Discoverer was used to determine phosphosite localization confidence. To match *ptm*RS scores to Progenesis MASCOT output files, two rounds of Excel vlookup functions were used: 1) to retrieve scan numbers from the TOP hit per feature number, and 2) to retrieve *ptm*RS score using the corresponding scan number. ‘Normalyzer’ software [32] was used to assess the suitability of various types of normalization strategies on the data. Amongst normalization strategies, VSN-G (group-based variance stabilizing normalisation) performed well under various quality control metrics indicating a reduction in technical variance. The ‘vsn’ R/Bioconductor package [33] was then used to normalize log transformed peptide abundance data for the subsequent stages of analysis. Pairwise t-test and cross condition ANOVA statistical testing, followed by Benjamini-Hochberg global correction was subsequently also performed in R for any replicate groups without missing values. Unless otherwise described, all plots were generated in R.

### Network analysis

All proteins with quantified phosphopeptides in 2 or more repeats of any condition were combined for network analysis of RelA-associated proteins. Common contaminants that bind non-specifically in IP experiments were removed by filtering protein accessions against a CRAPome database (https://www.crapome.org) [34] of contaminants observed with Streptactin-HA IPs from U2OS cells (CC405, CC406 and CC410). All proteins not present in the contaminant database were entered into STRING (https://string-db.org, v11.0) to generate a network of protein associations [35]. The generated network was filtered to contain only high confidence associations (interaction score ≥0.7) with experimental or database evidence only. Proteins with no high confidence interactions were removed from the network and the remaining nodes were clustered using the Kmeans algorithm (K=9). The resulting network was imported into Cytoscape [36] with nodes recoloured, grouped into the top 9 Kmeans clusters and node shapes for proteins unique to HU or ETO changed to squares or diamonds respectively. For the top 9 Kmeans clusters, nodes corresponding to proteins with differentially regulated phosphopeptides in response to either HU, ETO or both treatments, with respect to the untreated control, were outlined in red, blue or black respectively.

### Functional enrichment analysis

All functional enrichment analysis was performed using the functional annotation tool in DAVID (https://david.ncifcrf.gov, v6.8) against a background of the human proteome [37,38]. For biological process GO term pie charts, all significantly enriched (Benjamini adj.*p*-value ≤ 0.05) GO biological process terms, with associated Benjamini adj.*p*-values, were summarised using REVIGO [39] then visualised with the CirGO Python package [40]. Bubble plots including significantly enriched (Benjamini adj.*p*-value ≤ 0.05) GO biological process (BP), molecular function (MF), cellular component (CC) and overrepresented keywords, filtered to remove duplicate terms, were plotted in R.

### Kinase Prediction/Sequence motif analysis

Kinase-substrate prediction for all differentially regulated (*t*-test, *p* ≤ 0.05) phosphosites was performed using NetPhorest [41,42]. Briefly, for all ‘class I’ phosphorylation sites (*ptm*RS ≥0.75) protein identifiers together with the position of the phosphorylated residue were submitted to NetPhorest using the high-throughput web interface, with the top scoring prediction for each site reported. Kinase predictions were summarised for all up- or down-regulated phosphosites with the resulting data plotted in R. For sequence motif analysis, sequence windows encompassing 7 amino acids either side of the phosphorylated residue were extracted for all phosphosites significantly up- or down-regulated (*t*-test, *p* ≤ 0.05) with respect to the untreated control. Consensus sequence motifs were generated with iceLogo v1.2 [43] against a background of the precompiled human Swiss-Prot composition using percent difference as the scoring system and a *p*-value cut-off of 0.05.

## RESULTS

### Selection of DNA damage timepoints

To identify changes in the phosphorylation status of RelA and its protein binding partners in response to different types of DNA damage, we first examined the kinetics of DNA damage after treatment with either the DSB inducer ETO, or the replicative stress inducer HU in the HA-RelA U2OS cell line. Using gamma-H2AX as a marker [44,45], we observed rapid DNA damage (within 30 min) following treatment with either ETO or HU (Fig. 1; Supplementary Fig. 1). Gamma-H2AX levels peaked faster with ETO than HU, but sustained (and maximal) levels were observed with both agents following ∼ 2 hours of continuous treatment. A time-course of the effect of these different DNA damaging agents on the RelA phosphorylation interactome was therefore evaluated by treatment of the HA-RelA U2OS cells with either ETO or HU for 0, 30, 60 or 120 minutes prior to immunoprecipitation (IP) of the HA-tagged RelA. Immunoprecipitated proteins were subject to titanium dioxide (TiO_2_)-based phosphopeptide enrichment prior to LC-MS/MS with label-free peptide quantification (Fig. 1).

**Figure 1.**
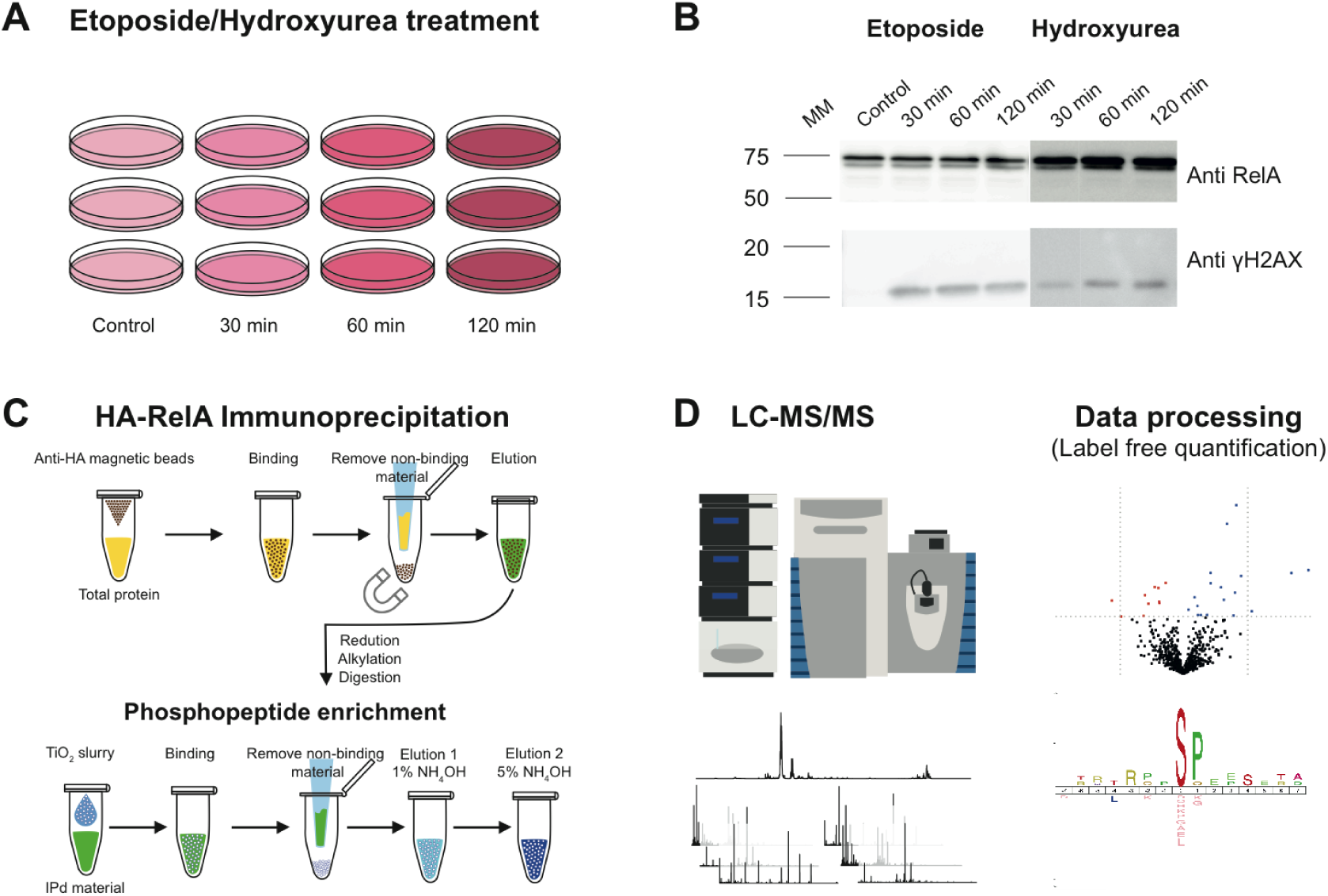
Workflow for evaluation of DNA damage response of HA-RelA U2OS cells with either Etoposide (ETO) or Hydroxyurea (HU). (**A**) HA-RelA U2OS cells were treated with either 0.1% DMSO (control), ETO (50 μM) or HU (2 mM) for the times indicated to induce DNA damage. (**B**) Following treatment with either ETO or HU, HA-RelA U2OS cell lysates were immunoblotted with antibodies against either γH2AX as a marker of DNA damage, or RelA/p65. (**C**) Workflow for evaluation of the phosphorylated RelA interactome. Cells were treated as in (**A**), and RelA-bound proteins immunoprecipitated with an anti-HA antibody. Proteins were subjected to tryptic digestion and TiO_2_-based phosphopeptide enrichment. (**D**) Peptides were analysed by LC-MS/MS using a QExactive HF and subjected to label-free quantification following appropriate normalisation.

**Supplementary Figure 1.**
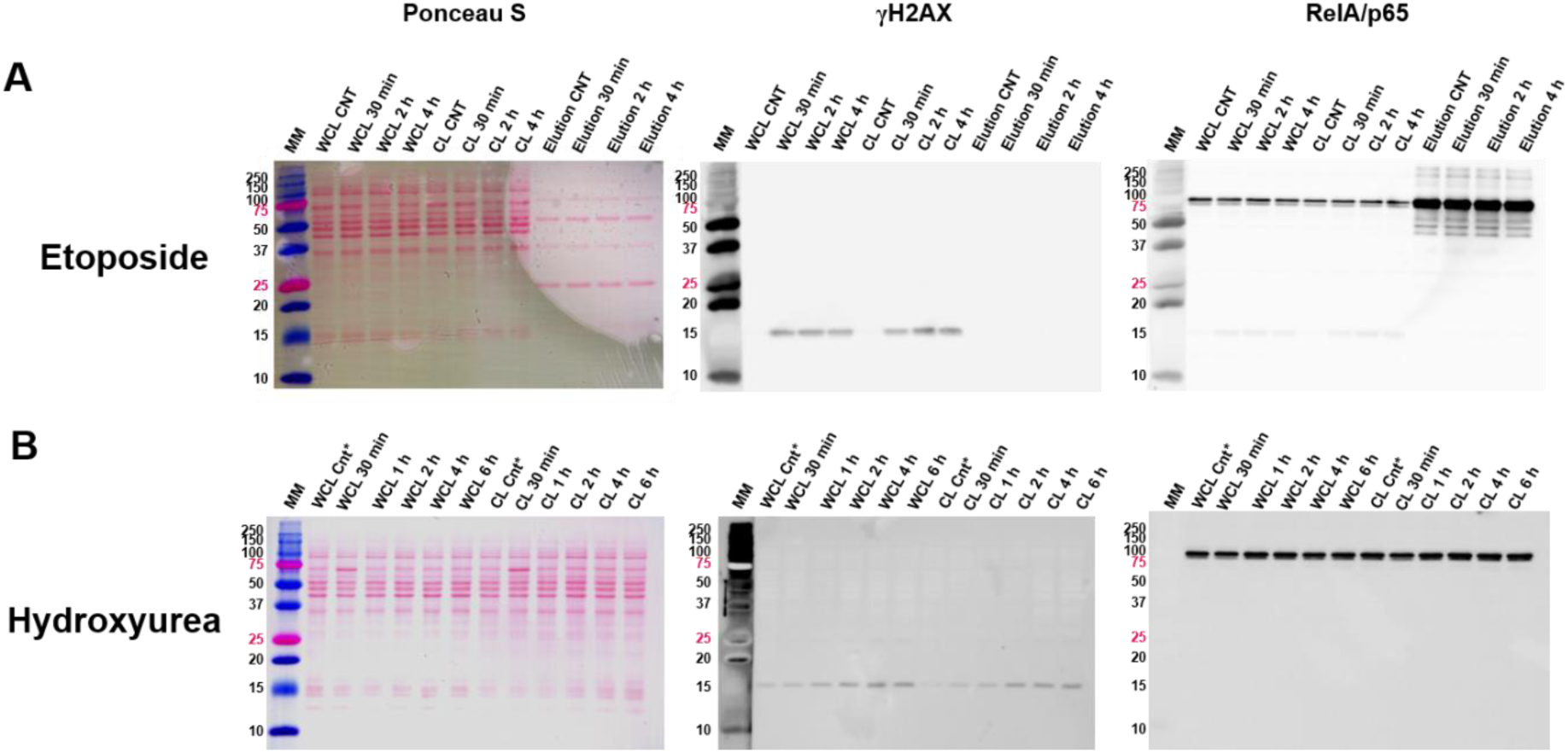
Time course of treatment of HA-RelA U2OS cells with etoposide or hydroxyurea. HA-RelA U2OS cells were treated for the specified times with either etoposide (A) or hydroxyurea (B) as indicated to assess DNA damage. Cell lysates were separate by SDS-PAGE; total protein was visualised with Ponceau S staining. Gels were then subjected to immunoblotting with antibodies against either γH2AX as a marker of DNA damage, or RelA/p65. MM: molecular markers; WCL: whole cell lysate; CL: cleared lysate; Elution: eluant from anti-HA beads; CNT: control. Control cells for hydroxyurea treatment were exposed to DMSO for 6 h to match longest treatment time.

### Network of RelA-associated proteins

Using the identified (phospho)peptides as proxy for protein identifications, the list of RelA bound proteins under all conditions was filtered against a ‘CRAPome’ database of contaminants commonly observed in Streptactin-HA IPs from U2OS cells (CC405, CC406 and CC410), which provided the closest match to the experimental workflow described here. Phosphopeptide quantification data was normalised and subjected to ANOVA testing across all conditions, and individual t-tests across all pairs of conditions. Using this list, the temporal effect of treatment with either ETO or HU on RelA bound phosphoproteins was evaluated for all those proteins quantified in at least 2 biological replicates, under a given condition.

To generate a network of high confidence interactors, the remaining 815 RelA-associated proteins combined across all conditions (Supp. Table 1) were inputted into STRING (v11.0) [35] using experimental and database sources only, and clustered using the Kmeans algorithm (k=9). Twelve of these proteins failed to yield high confidence interactions and therefore were filtered out of subsequent datasets (Fig. 2). Gene Ontology (GO) enrichment analysis of biological function for these 803 proteins within the top 9 network clusters confirmed association of RelA with a number of IκB kinase/NF-κB signalling proteins (Fig. 2A), including RelB, c-Rel, p105/p50, p100/p52, IκBα, IκBβ, and the DNA damage response (Fig. 2B). Clusters of proteins across a number of core cellular processes were also identified in the RelA interactome, including key proteins involved in cell division, transcriptional regulation, mRNA/rRNA processing and MAPK/ERK/VEGF signalling (Fig. 2 [A-H]). Amongst these interactions partners, we also confirmed observation of a number of known RelA binding proteins, including *e.g*. replication factor C (RFC1), p300, SP1 and HDAC6 [46–49].

**Figure 2.**
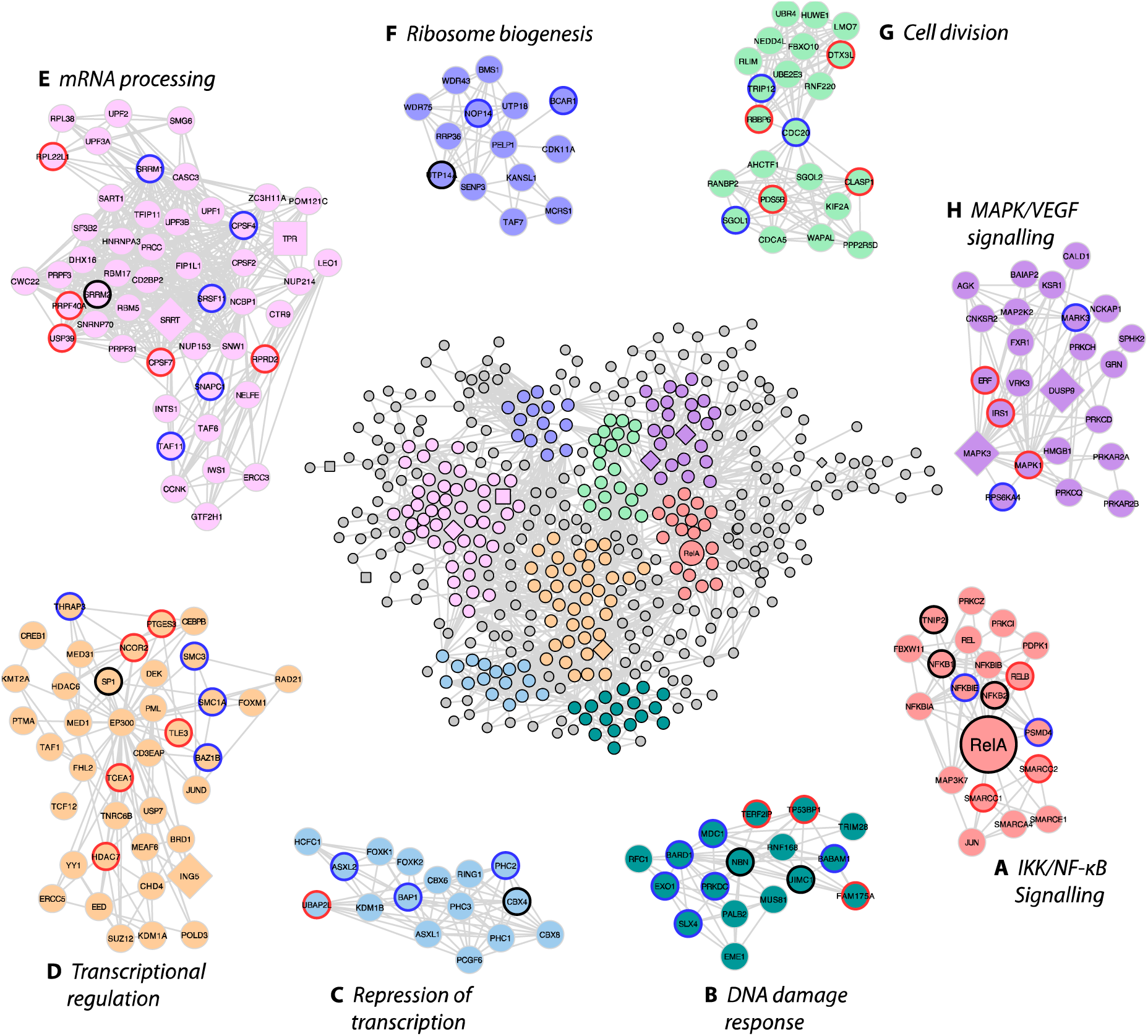
Network map of RelA interacting phosphoproteins. Proteins from phosphopeptides quantified following HA-RelA immunoprecipitation in control (0.1% DMSO control), ETO (50 μM) or HU (2 mM) treated U2OS cells were evaluated with STRING (v11.0) [35] for prior experimental evidence of interaction. Kmeans clustering was used to identify the top 9 enriched GO clusters of biological function and plotted using Cytoscape [36]. The shape of the nodes represent the conditions under which they was identified: round nodes represents proteins identified across all conditions; diamond nodes indicate proteins unique to ETO treatment; square nodes indicate proteins unique to HU treatment. (**A**) red nodes represent those proteins mapped to IKK/NF-κB signalling; (**B**) dark green nodes represent those proteins mapped to DNA damage response/DNA repair; (**C**) blue nodes represent those proteins mapped to repression of gene expression; (**D**) yellow nodes represent those proteins mapped to transcriptional regulation; (**E**) pink nodes represent those proteins mapped to mRNA processing/splicing; (**F**) violet nodes represent those proteins mapped to ribosome biogenesis/rRNA processing; (**G**) light green nodes represent those proteins mapped to cell division/mitosis; (**H**) purple nodes represent those proteins mapped to MAPK/VEGF signalling. Proteins outside of these 8 enriched clusters have nodes in grey. The outline colour of the nodes within the 8 clusters shows proteins with differentially regulated phosphopeptides following treatment with either ETO (blue), HU (red), or both (black).

To determine the temporal response in the RelA interactome of phosphorylated proteins to the different DNA damaging agents, we evaluated those quantified proteins exhibiting a statistically significant (t-test, *p* ≤ 0.05) change in response to time-dependent treatment with ETO or HU, in comparison to control (DMSO treated) cells. Interestingly, the constituents of the RelA interactome changed very little in response to DNA damage, with the vast majority of the phosphoproteins identified being observed before and after exposure to either ETO or HU (round nodes, Fig. 2). Of the 803 protein binding partners with high confidence interactors, only 7 were unique to ETO (diamond nodes, Fig. 2; Supp. Table 2), while 8 were specific to HU (square nodes, Fig. 2; Supp. Table 3). Of the 7 ETO-specific proteins identified in this network, 3 are essential components of the ERK/MAPK signalling pathway: the protein kinases ERK1 and PAK, and DUSP9, a dual specificity phosphatase that preferentially targets the MAPK/ERK family. A fourth member of this ETO-specific cohort, AAED1 (which has no known common confident interactors and is therefore not represented in Fig. 2), is also reported to positively regulate the ERK/MAPK (and AKT1) pathway, ultimately leading to upregulation of hypoxia-inducible factor (HIF)-1α and enhanced glycolysis [50]. In contrast, the 8 HU-specific proteins exhibited diverse biological functions, with only one of these proteins, nucleoprotein TPR, falling into one of the top 8 network clusters (Fig. 2A).

Also highlighted in the nodes of the top 8 network clusters are the proteins for which we identified statistically significantly differentially regulated phosphopeptides (t-test, *p* ≤ 0.05) in response to treatment with either ETO or HU (or both) with respect to the (DMSO treated) control cells (Fig. 2 [A-H]; Supp. Table 3). Interestingly, the proteins with differentially regulated phosphopeptides following either ETO or HU treatment are largely distinct between the two types of DNA damaging agents, and are interspersed across the network clusters rather than being specific to certain functional biological groups (Supplemental Figure 2-gifs). These data thus suggest that whilst there was relatively little change in the RelA-associated phosphoproteins as a function of DNA damage, the phosphorylation states of those associated proteins was dependent on both the type and duration of the induced DNA damage response.

### RelA is minimally differentially phosphorylated in U2OS cells in response to ETO or HU treatment

Looking specifically at RelA, we observed good sequence coverage of the N-terminal region (>70%; 46% overall; Supp. Fig. 3), allowing us to quantify most of the known phosphorylation sites in the *N*-terminal half of the protein including; Ser42, Ser45, Ser131, Ser203/Ser205, Ser238, Ser240 and S316. Two novel phosphorylation sites at Thr54 and Ser169 were also observed (Supp. Fig. 3). No phosphorylation sites were confidently identified in the *C-*terminus of RelA containing the transactivation domains (TAs), in part due to the limited ability to generate suitable tryptic peptides for analysis (as documented previously [26]). Although phosphopeptides covering the extreme C-terminus (from residue 503 and including the known sites of phosphorylation at T505, S529 and S536) were observed, the site of phosphorylation could not be unambiguously defined.

**Figure 3.**
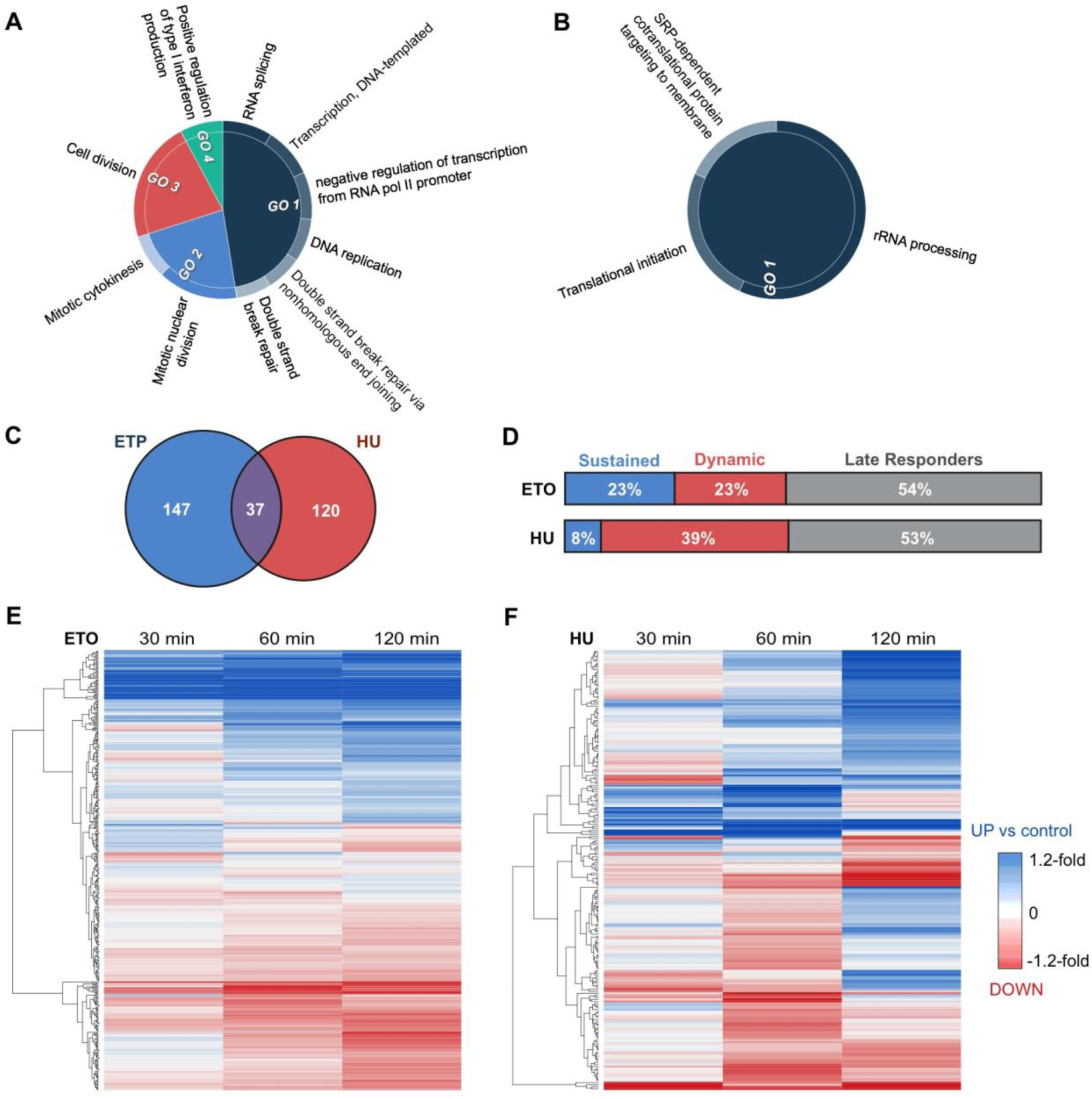
Etoposide and Hydroxyurea induced different phosphorylation dynamics in the RelA network. GO term biological process enrichment was examined for all phosphopeptides exhibiting a statistically significant change in response to either (**A**) etoposide (ETO) or (**B**) hydroxyurea (HU). (**C**) Overlap of those proteins with significant change in phosphopeptide abundance at one or more time points with ETO or HU. (**D**) Distribution of phosphopeptide changes between sustained, dynamic and late responders for ETO vs HU treatment, where ‘Late Responders’ indicates a change at 120 minutes only, ‘Sustained’ indicates a change at 30 or 60 minutes which is sustained throughout, and ‘Dynamic’ indicates a change at 30 or 60 minutes which is reversed. (**E, F**) Heatmaps of the fold change in phosphopeptide levels following treatment with either ETO (**E**) or HU (**F**) at 30, 60 or 120 min relative to control levels. Hierarchical row clustering was performed on the log2 fold changes.

Overall, the changes observed in RelA phosphorylation in response to DNA damage under these conditions were very limited. Only Ser131 was statistically significantly regulated by both the DSB-inducing ETO and HU: pSer131 increased marginally by 1.03-fold (with respect to the DMSO-treated control) at 120 min (*p-*value = 0.047) in response to ETO, responding much more rapidly to HU, increasing 1.05 fold by 30 min (*p-*value = 0.046), decreasing slightly (although not statistically) by 60 min, and returning to baseline levels by 120 min. We have previously shown that this site is also responsive to TNFα treatment of U2OS cells, and phosphorylated *in vitro* by IKKβ, in a manner that is significantly enhanced in the presence of the RelA dimerization partner, p50, and reduced in the presence of IκBα [26]. In contrast, Ser45, a phosphorylation site that we previously demonstrated plays a critical role in reducing the ability of RelA to bind to DNA (alongside Ser42, which was not observed in the HU-treated cells) [26], was reduced by 1.1 fold (*p-*value < 0.01) 60 min after cellular treatment with HU, suggesting an HU-mediated regulation of RelA transcription, in part through Ser45 phosphorylation (Supp. Fig. 3).

**Supplementary Figure 3.**
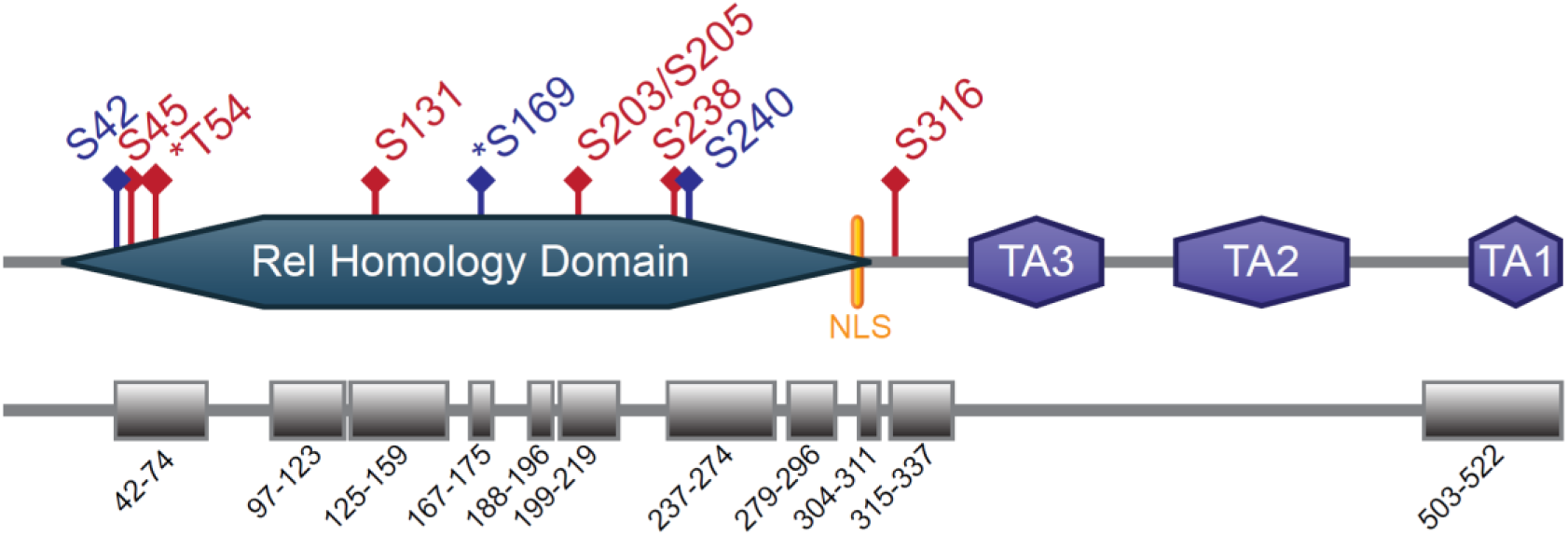
Schematic of RelA protein coverage. Domains and quantified sites of phosphorylation are indicated. Blue phosphorylation sites were quantified in ETO only, red were quantified in both ETO and HU. Novel phosphorylation sites are represented with an asterix. Protein sequence coverage is represented by the boxes (and amino acid residue numbers) in the bottom panel. TA = transactivation domain; NLS – nuclear localisation sequence.

### ETO and HU induced different phosphorylation dynamics in the RelA network

Of the 696 proteins identified in the RelA network in the presence of ETO, 184 exhibited ETO-dependent changes in phosphorylation status, with a total of 312 phosphopeptides exhibiting a statistically significant change. GO term analysis of biological processes of these 184 proteins revealed enrichment primarily of proteins involved in transcription (*p* < 0.001), RNA processing (*p* < 0.001), cell division (*p* < 0.001) and, unsurprisingly, the DNA damage response (*p* < 0.001) (Fig. 3A). Slightly fewer changes arose in response to HU, with 235 phosphopeptides from 157 proteins exhibiting a statistically significantly change compared to control (Fig. 3B), those primarily being involved in translation (*p* < 0.001) and rRNA processing (*p* < 0.001). Interestingly, only 37 of the proteins exhibiting dynamic changes in response to treatment were common between the two types of DNA damaging agent, indicating significant differences in the mechanisms whereby RelA mediates the outputs from these two DNA damaging agents (Fig. 3C).

We next evaluated the temporal dynamics of these phosphorylation changes as a function of treatment type (Fig. 3D). Intriguingly, ETO and HU induced very different dynamics in phosphosite regulation. Indeed, approximately one-quarter (∼70 phosphopeptides) of the ETO-regulated phosphopeptides, were responsive within 60 min of treatment, with the change in phosphorylation status being maintained over the duration of the 120 min experiment (Fig. 3D, E). In contrast, only 8% (∼20 phosphopeptides) of HU-dependent phosphorylation changes occurred within the first 60 min and were then sustained for the duration (Fig 3D, F). Overall, HU-mediated phosphorylation changes were much more dynamic, with 39% of the phosphopeptides quantified in these experiments significantly regulated (being observed at either increased, or reduced levels), before reverting back, either to control levels, or in some instances, in the opposite direction. Just over half of those phosphopeptides that were differentially regulated by either ETO or HU did not exhibit a statistically significant change until after more than 60 min of continuous treatment, *i.e*. they were late responders that were only observed in these experiments 120 min after exposure (Fig. 3D).

### Phosphorylation of RelA-associated proteins in response to etoposide

Of the 312 phosphopeptides that were differentially regulated by ETO (with respect to the untreated control (*p* ≤ 0.05)) across all time-points, 151 increased in abundance, while 159 were down-regulated (Supp. Table 2). Only two phosphorylation sites exhibited significant temporal dynamics (with levels being significantly elevated or decreased at sequential time-points), although the selective nature of the timing at which these measurements were made could be masking additional dynamics. Overall, the relative fold change in phosphopeptide abundance (up- or down-) was relatively small, with the greatest statistically significant fold change being ∼1.4 (Fig. 4A-C; Supp Table 2).

**Figure 4.**
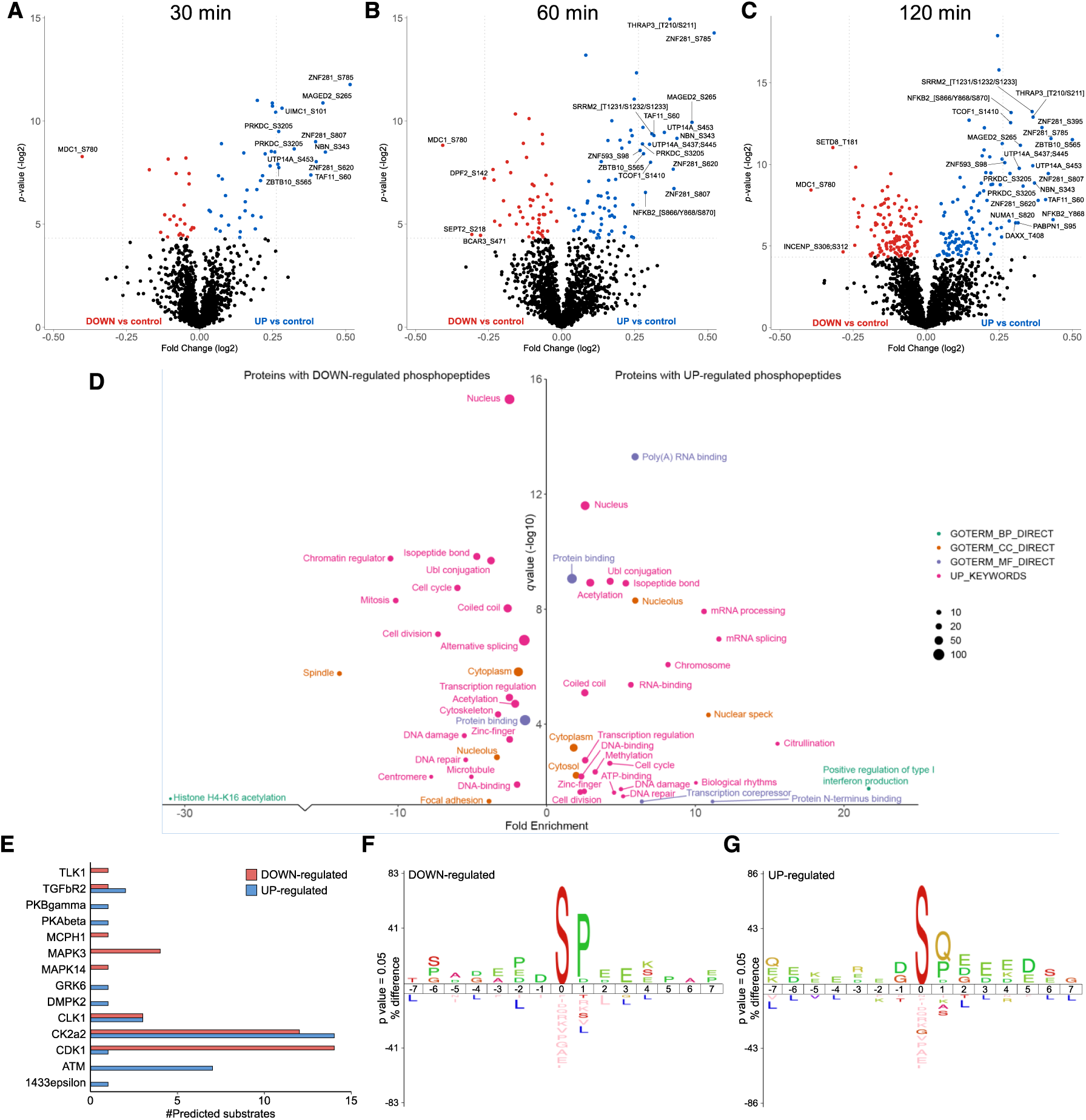
Etoposide-mediated phosphopeptides changes in the RelA network. Volcano plots showing fold changes in phosphopeptide abundance following Bayesian statistical analysis to evaluate significant differences as a function of (**A**) 30 min, (**B**) 60 min or (**C**) 120 min treatment with ETO. Log_2_-fold change are presented as a function of the –Log_2_ p-value; differentially down-regulated (red) or up-regulated (blue) phosphopeptides with a p-value ≤0.05 are highlighted. Select data points are annotated with their protein accession number and site of phosphorylation. (**D**) GO term enrichment analysis using DAVID of proteins with significantly regulated phosphopeptides in response to ETO (all time points) relative to control. Phosphopeptides with a Benjamini-Hochberg adjusted p-value ≤0.05 are labelled. BP = biological process (green); CC = cellular compartment (yellow); MF = molecular function (purple); UP = UniProt keyword (pink). (**E**) NetPhorest kinase-substrate prediction for significantly down- (red) or up-regulated (blue) phosphosites. IceLogo sequence analysis of (**F**) down-regulated or (**G**) up-regulated phosphosites.

In-depth functional enrichment analysis of proteins that were either up- or down-regulated following ETO exposure revealed some key differences in terms of the biological functions that were regulated (Fig. 4D). RelA-associated proteins with an ETO-mediated reduction in phosphopeptide levels showed significant enrichment for GO terms including mitosis, chromatin regulation (including H4-K16 acetylation), transcriptional regulation and DNA damage repair (Fig. 4D). pSer780 on MDC1, the Mediator of DNA damage checkpoint protein 1, exhibited the greatest down-regulation of all phosphosites in response to ETO across the time-points investigated (1.3-fold down; *p* = 1.95E-04) (Supp. Table 2, Fig. 4A-C). This phosphorylation site has previously been identified has being induced in response to ionising radiation (IR) [51] and ultraviolet (UV) radiation [52], both of which induce DSB, although the physiological function of this phosphorylation site has not yet been defined. We show here that the proportion of Ser780 phosphorylated MDC1 that contributes to the RelA interactome decreases in response to ETO, suggesting that DNA-damage induced pSer780 may serve to disrupt the interaction of MDC1 with RelA transcriptional complexes.

It is interesting to note that many more phosphopeptides were significantly upregulated than were decreased in the RelA interactome in response to ETO, particularly in the later time points. Transcriptional regulation and DNA damage repair were also among the GO terms enriched for those proteins with up-regulated phosphorylation sites. However, there was only an ∼8% overlap in common proteins with differentially regulated (up or down in response to ETO) phosphopeptides (Supp. Table 2), suggesting co-ordinated regulation of these key processes largely through different protein effectors rather than differential phosphorylation-mediated regulation of the same proteins. Also among the GO terms enriched in the list of proteins with up-regulated phosphopeptides in response to ETO were mRNA processing, ATP binding, transcription corepressor functionality and type I interferon production, including the NF-κB signalling proteins (RelA (pSer131), NFKB1 (p105/p50) (pSer923;Ser927) and NFKB2 (p100/p52) (pSer858, pTyr868 and pSer870), with pSer619 on the RNA helicase DDX41 and pSer3205 on the DNA-dependent protein kinase PRKDC (Fig. 4D). Multiple phosphopeptides from the transcription factor zinc finger protein 281 (ZNF281/GZP1/ZBP99), containing pSer395 (*p* = 2.1E-02), pSer620 (*p* = 1.1E-04), pSer785 (*p* = 1.9E-06) and pSer807 (*p* = 1.7E-03), were significantly upregulated across all time points (between 1.2-1.4-fold) in the RelA interactome in response to ETO, but not in response to HU. ZNF281 has recently been reported to be recruited to DSBs, following interaction of its zinc finger domain with XXCR4 [53], where is plays an important role in non-homologous end-joining upon DNA damage, and consequently the maintenance of cell viability. Supporting our findings of increased phosphorylation in response to DSB, these authors also reported a slight reduction in the recruitment of a non-phosphorylatable S785A/S807A ZNF281mutant to sites of DNA damage. Based on our data, we would suggest that phosphorylation of ZNF281 on Ser395 and Ser620 may also be required for its optimal recruitment to DNA lesions and subsequent DNA repair. In an attempt to understand the signalling pathways and possible kinase-substrate relationships feeding into the ETO-dependent changes in phosphorylation of the RelA-binding partners, the ‘class I’ localized phosphosites (*ptm*RS ≥0.75) with *p*-value ≤ 0.05 were used as input sequences for the NetPhorest kinase-substrate prediction algorithm [34,35] (Fig. 4E). Of the 117 down-regulated and 107 up-regulated phosphorylation sites that passed the above thresholds, NetPhorest gave kinase-substrate predictions for 37 and 32 sites respectively. Among the down-regulated sites, CDK1 had the highest number of predicted substrates (14, 38%) which accounts, at least partially, for the notable enrichment across all the ETO down-regulated phosphorylation sites for Pro in the +1 position with respect to the site of phosphorylation [54] (Fig. 4F). The identification of four predicted substrates for ERK1 (MAPK3), which is also a Pro-directed protein kinase [55], likely also contributes to the identified sequence motif, and correlates with our observations of this enzyme, and importantly, a cognate phosphatase, DUSP9, in the RelA network under these conditions.

IceLogo sequence analysis of the ETO up-regulated phosphosites showed significant enrichment for Gln (and to a lesser extent Pro) at the +1 position with respect to the site of phosphorylation (Fig. 4G). Positions +2 to +5 also exhibited a strong preference for acidic residues (Asp/Glu), consistent with observation of 14 predicted substrates for casein kinase II alpha (CK2α2) in the NetPhorest output (Fig. 4E). Perhaps not surprisingly given its somewhat promiscuous substrate repertoire, and an involvement in diverse cellular processes (including regulation of numerous transcription factors such as NF-κB), CK2α2 was also predicted as the regulatory enzyme for a number of the significantly down-regulated phosphorylation sites. The striking observation of Gln at +1 in 36 of the up-regulated phosphorylation sites (Fig. 4G) is in agreement with previous reports of ATM/ATR-dependent regulation of the RelA NF-κB pathway in response to DSB (albeit following cellular exposure to TNFα rather than etoposide) [56]. ATM/ATR kinases are well known to preferentially phosphorylate Ser/Thr residues following by a Gln, with substrates often containing several closely spaced SQ/TQ motifs in regions termed SQ/TQ cluster domains (SCDs) [57]. Whilst just 7 of the 36 SQ motif-containing sequences were predicted by NetPhorest to be ATM sites, it is possible that many more of the differentially regulated phosphosites that match to this consensus are potential ATM/ATR substrates, given the limitations of prediction software. While phosphorylation at any given site can be induced by more than one protein kinase in cells, the NetPhorest algorithm is best used as a guide and will only assign a single potential enzyme, based on the best fit, and is therefore not fully representative of cellular possibilities.

### Phosphorylation of RelA-associated proteins in response to hydroxyurea

Upon HU treatment, 235 phosphopeptides from 157 proteins in the RelA interactome network were significantly differentially regulated with respect to the DMSO treated control (*p* ≤ 0.05; Fig. 5 A-C). Of those, 121 phosphopeptides increased in abundance, while levels reduced for 109 phosphopeptides. Among the phosphorylation sites exhibiting a dynamic response over the time-course of HU treatment, five showed a bi-directional response, increasing in some time points, but reducing in others, with respect to control levels (Supp. Table 3). RelA-associated proteins with statistically significantly reduced phosphopeptide levels in response to HU-mediated DNA damage were notably enriched for GO terms encompassing ribosome biogenesis, mRNA processing and translation (Fig. 5D). Notably, whilst translation was enriched among the proteins effected by HU treatment there was no significant enrichment for proteins involved in translation with ETO. In contrast, elevated phosphopeptide levels were observed for processes such as chromatin regulation and chromosomal rearrangement, and to a lesser extent, transcriptional regulation and mitosis (Fig. 5D). Whilst transcriptional regulation was among the GO terms enriched in response to HU, it was only marginally enriched, and with a much lower significance than our observations with ETO, suggestion a more much limited effect on transcriptional regulation with HU than with ETO (Fig. 4D, Fig. 5D).

**Figure 5.**
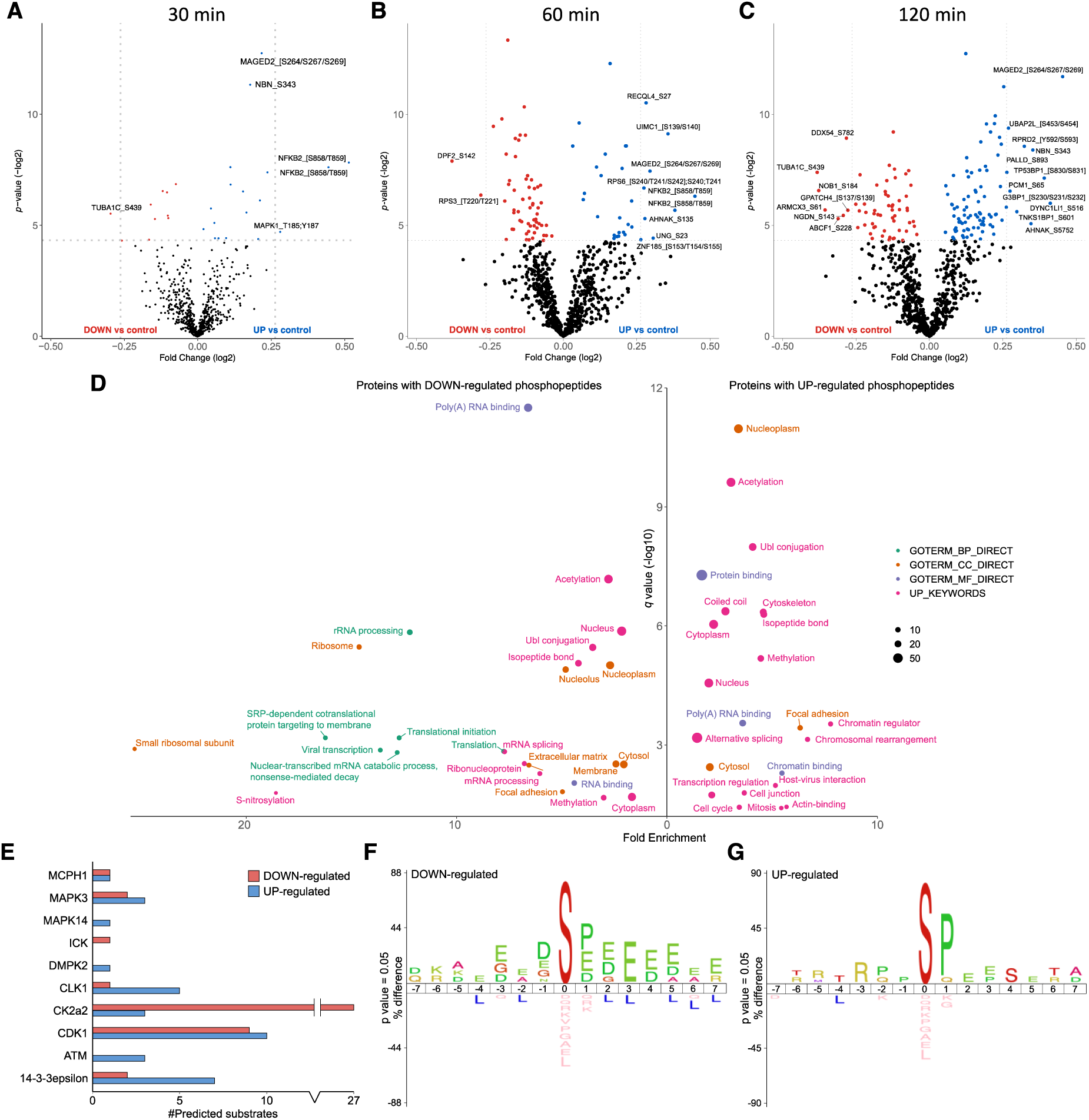
Hydroxyurea-mediated phosphopeptides changes in the RelA network. Volcano plots showing fold changes in phosphopeptide abundance following Bayesian statistical analysis to evaluate significant differences as a function of (**A**) 30 min, (**B**) 60 min or (**C**) 120 min treatment with HU. Log_2_-fold change are presented as a function of the –Log_2_ p-value; differentially down-regulated (red) or up-regulated (blue) phosphopeptides with a p-value ≤0.05 are highlighted. Select data points are annotated with their protein accession number and site of phosphorylation. (**D**) GO term enrichment analysis using DAVID of proteins with significantly regulated phosphopeptides in response to HU (all time points) relative to control. Phosphopeptides with a Benjamini-Hochberg adjusted p-value ≤0.05 are labelled. BP = biological process (green); CC = cellular compartment (yellow); MF = molecular function (purple); UP = UniProt keyword (pink). (**E**) NetPhorest kinase-substrate prediction for significantly down- (red) or up-regulated (blue) phosphosites. IceLogo sequence analysis of (**F**) down-regulated or (**G**) up-regulated phosphosites.

In agreement with the γH2Ax immunoblotting data (Fig. 1B) which indicated a slower cellular response to HU than ETO (see the heatmaps in Fig. 3F), very few significant changes in phosphopeptide abundance were observed 30 min after treatment with HU (Fig. 5A). Interestingly, the NFKB2 (p100) peptide phosphorylated on either Ser858/Thr859 (site ambiguous) exhibited the greatest increase of any phosphopeptides at this time point, with a 1.4-fold change (Supp. Table 3; being observed in two charge states) (Fig. 5B). This increase in HU-mediated phosphorylation is supported by additional quantification of a smaller peptide in which the site of phosphorylation was defined as Ser858 (Supp. Table 3), a site which was also significantly upregulated following exposure to ETO (Supp. Table 2). Of note, these peptides (and others in the C-terminus of p100 that are differentially regulated in response to treatment) are specific to p100, rather than the transcriptionally active p52 cleavage product, suggesting DNA damage-mediated regulation of either p100 processing, or its ability to bind (directly or indirectly) to RelA.

The second most differentially elevated phosphopeptide after 30 (and 60) min treatment with HU was the doubly phosphorylated pThr185/pTyr187-containing peptide from ERK2. This phosphopeptide lies in the kinase activation loop and phosphorylation of both of these sites is required for catalytic activation of ERK2 [58,59]. An early consequence of HU treatment on the RelA interactome network thus appears to be activation of bound ERK2, which contrasts with our observations with ETO, where there were no significant changes induced in MAPK1/ERK2 phosphopeptides. Together, these findings are curious in light of the fact that the C-terminal region of p100 is reported to bind to inactive ERK2, thereby preventing its phosphorylation and nuclear translocation. The fact that we observe increased phosphorylation of ERK2 alongside phosphorylation in the C-terminal ‘death domain’ of p100 may suggest a role for these p100 phosphorylation sites in regulation its ability to interact with ERK2 [60].

Nibrin, a member of the MRN complex which plays a critical role in DSB repair and cell cycle checkpoint control, also exhibited elevated Ser343 phosphorylation by 30 min HU treatment. Interestingly, the pSer343 site observed falls within an SQ motif which has previously been shown to be phosphorylated by ATM in response to IR, and is believed to play a role in intra-S phase checkpoint activation [61,62] (Fig. 5A and 5E). This same phosphorylation site was also rapidly elevated in response to ETO (Fig. 4B), confirming its apparent involvement in a generic ATM-mediated DNA damage response [61,62]. By 60 min, we also observed reduced levels of pSer142 on DPF2 (Zinc finger protein ubi-d4), a known regulator of the non-canonical NF-κB pathway [63], and pThr220/221 (site ambiguous) on the RelA binding partner RPS3 (40S ribosomal protein S3), both of which have previously been reported to be elevated in HEK293 cells following 3 h UV exposure [52]. Although binding of RSP3 to RelA is thought to enhance its ability to bind DNA *in vitro* [64], RSP3 functions as a negative regulator of H_2_O_2_-mediated DNA repair [65]. It is possible therefore that pThr220/221 on RPS3, and pSer142 on DPF2, serve to modulate the RelA transcriptional complexes and/or the consensus promoter regions for NF-κB binding, that are required for DNA repair in response to HU.

Using NetPhorest [41,42], we were able to predict possible kinase-substrate relationships for 46% (77 out of 167) of the ‘class I’ phosphosites (*ptm*RS ≥0.75) that were regulated in response to HU (Fig. 5E). CK2α2 was predicted as the kinase responsible for the vast majority (27 out of 43) of the phosphosites reduced in response to HU, based on the prevalence of acidic residues C-terminal to the site of phosphorylation (in particular at +3) clearly evident in the sequence analysis (Fig. 5F). Consequently, HU might have significant effects on CK2α2 (or other acidophilic kinases) that feed into the RelA-mediated DNA damage response. Prevalent among both up-and down-regulated phosphosites were predicted substrates of cyclin dependent kinase 1 (CDK1) which, together with the prediction of substrates for ERK1 (MAPK3) and p38α (MAPK14), contributes to the enrichment of Pro at the +1 position in both the HU up- and down-regulated phosphosites (Fig. 5F and 4G). Also among the up-regulated phosphopeptides are a number of predicted ATM substrates which, whilst only representing a small proportion of the total number of HU-regulated phosphosites, corresponds with the small enrichment for Gln in the +1 position among the up-regulated phosphosites. (Fig. 5E and 5G).

Comparing the putative kinase-mediated regulation of substrates for those differentially regulated phosphopeptides in the RelA interactome in response to the two different types of DNA damaging agents revealed some interesting observations: although ATM-predicted substrates were enhanced with both ETO and HU, over twice as many putative substrates were elevated with ETO. Predicted substrates for CK2α2 are prevalent in the differentially regulated (both up- and down-) phosphosites under both conditions, although HU induces a far greater reduction in levels of putative CK2α2 substrates than ETO. In contrast, putative substrates for the dual specificity protein kinase CLK1, which has a role in regulating alternative splicing, are notably elevated in response to HU (Fig. 5E), but not particularly with ETO (Fig. 4E). Interestingly, binding sites for 14-3-3ε, which generally conform to the RXXpS/TXP consensus, were elevated under both conditions, but particularly in response to HU. 14-3-3 proteins are known to be involved in the cellular response to DNA damage; 14-3-3ε in particular, modulates NF-κB signalling in part by virtue of its ability to bind phosphorylated TAK1 (and its cognate phosphatase PPM1B), and thereby regulate its anti-apoptotic capabilities [66,67]. In response to DNA damage stress, this manifests as inhibition of the anti-apoptotic activity of TAK1, promoting an apoptotic phenotype that is enhanced by the 14-3-3ε-mediated disruption of DP-2 from the E2F transcriptional complex [68]. The fact that we observe elevated putative 14-3-3ε binding sites following HU treatment (but significantly less so with ETO), is in agreement with the pro-apoptotic response of HU-treated cells. Consequently, we hypothesise that the ability of components of the RelA interactome to bind 14-3-3ε may be a driving factor in achieving the expected cell death/survival phenotype in response to these two DNA damaging agents.

## DISCUSSION

Protein phosphorylation is a key reversible, dynamic PTM that rapidly governs protein complex formation, particularly of transcription factor complexes, thereby coupling extracellular stressors with compensatory transcriptional output. To understand the role of phosphorylation in regulating the RelA transcriptional network in response to different types of DNA damage, we used a quantitative phosphoproteomics to explore the dynamics of the phosphorylated RelA interactome in response to either ETO or HU. Evaluating the RelA network under these conditions is pertinent, given the central role of this transcription factor in directing the cellular response towards either senescence or apoptosis. Exposure of U2OS cells to either ETO or HU had very limited effect on the phosphorylated RelA interactome; only 246 of the 815 RelA interacting phosphoproteins that we identified in total changed upon treatment. Of these, less than 1% were specific to the type of DNA stress, with 7 or 8 unique proteins (respectively) being identified in the RelA network following exposure of U2OS cells to either the DSB-inducing ETO or to HU. However, although there was little change in the make-up of this network, there was evidence of subtle, but significant, changes in the phosphorylation states of the RelA bound proteins as a function of both the type of and duration of the DNA damaging agent used. Not only did the anti-apoptotic agent ETO invoke a more rapid cellular response than HU, both in terms of maximal H2AX levels (Fig. 1B), and quantifiable changes in phosphopeptide levels (Figs. 3, 4 and 5; Supp. Table 2), but interestingly this study pointed to the differential regulation of key cellular processes and the involvement of unique signalling pathways in modulating the treatment-specific functions of RelA.

Examination of the RelA network hub (the bait) revealed two novel phosphorylation events amongst the RelA phosphopeptides that we were able to identify and quantify in this trypsin-based peptide analysis: Thr54 and Ser169, both of which are within the DNA binding domain. However, only Ser45 and Ser131 were statistically differentially regulated in response to either ETO or HU. While Ser131 was statistically elevated upon treatment with ETO at 120 min, the fold change was marginal (1.03-fold), with similar fold change (1.05-fold) being observed for this phosphosite in response to HU, albeit at a shorter 30 min time point. The RelA peptide containing pSer45 was unchanged upon ETO treatment, and was statistically downregulated by 60 min exposure to HU, again with low (1.05) fold change. Given that we were evaluating the total cellular pool of RelA, rather than specifically the nuclear portion where we would expect e.g. Ser45 phosphorylation to have a greater regulatory effect [26], it is perhaps not surprising that the phosphopeptides changes quantified in this study were relatively low.

Although a number of our findings of regulated phosphorylation site changes are in agreement with other published studies, the overall fold change in phosphopeptide levels across this study were relatively low (maximally 1.7 fold), and speaks to the necessity to avoid, wherever possible, implementation of a fold-change cut-off during this type of analysis. While ETO-regulated RelA bound phosphoproteins were primarily involved in transcription, cell division and mitosis, and DSB repair (as might be expected), HU-mediated changes were predominantly observed in proteins with functions in rRNA and mRNA processing, and translational initiation. Where changes were observed, ETO predominantly resulted in elevated phosphopeptide levels, with a number of the ETO-regulated phosphorylation sites identified having been reported previously to be modulated in response to either IR or UV radiation. It was interesting to note that transcriptional regulation and DNA damage repair proteins were enriched in those datasets with both elevated and reduced phosphopeptide levels, suggesting coordinated regulation of these processed through different effectors in response to DSB. In addition to the involvement in AMT/ATR signalling previously reported, potential substrates for which were much more prevalent following ETO treatment that HU, kinase substrate prediction suggested roles for CDK1 and ERK1 (or their cognate phosphatases), in regulating RelA complexes in response to ETO. In contrast, levels of the activation loop phosphopeptide in MAPK1/ERK2, traditionally thought to be a marker of kinase activity, were elevated in response to HU, but not ETO, suggesting differential MAPK isoform signalling in response to these two types of DNA damage. The prediction of substrates for p38α in the regulated phosphorylation sites following cellular exposure to HU (but not ETO), also points to a potential HU-specific regulation of this stress activated protein kinase. Finally our data point to a role for 14-3-3ε binding propensity in facilitating the cellular response to HU, which may be a contributing factor in differentiating the pro-and anti-apopotic phenotypes that are the result of prolonged cellular expose to these two type of DNA damaging agents, and we believe is worthy of further investigation.

## Author Contributions

CEE and NDP obtained funding. AC, CFF and CEE designed the experiments and analysed the data, with bioinformatics support from SP and ARJ. LS generated the HA-RelA U2OS cell line. CFF treated the cells, prepared the samples and did the immunoblotting studies. AC, CFF and PJB performed the MS analysis. AC and CEE wrote the paper with contributions from all authors, who also approved the final version prior to submission.

## Notes

The authors declare no competing financial interest. The mass spectrometry proteomics data have been deposited to the ProteomeXchange Consortium via the PRIDE partner repository with the dataset identifiers PXD019587 and PXD019589.

## Abbreviations

AGC: automatic gain control;
CID: collision-induced dissociation;
DSB: double strand break;
ETD: electron-transfer dissociation;
ETcaD: electron transfer with supplemental collision activation;
EThcD: electron-transfer higher energy collisional dissociation;
ETO: etoposide;
GO: gene ontology;
HCD: higher energy collisional dissociation;
HU: hydroxyurea;
IR: ionizing radiation;
MS: mass spectrometry;
PD: Proteome Discoverer;
PTM: post-translational modification;
RHD: Rel homology domain;
TD: transactivation domain;
UV: ultraviolet.
ATM: ataxia telangiectasia mutated;
CDK: cyclin-dependent kinase;
DMSO: dimethyl sulfoxide;
DSB: DNA double stranded breaks;
ERK: Extracellular signal-regulated kinase;
ETO: etoposide;
HU: hydroxyurea;
IR: ionising radiation;
MS: mass spectrometry;
MS/MS: tandem mass spectrometry;
PTM: post-translational modifications;
UV: ultraviolet

---

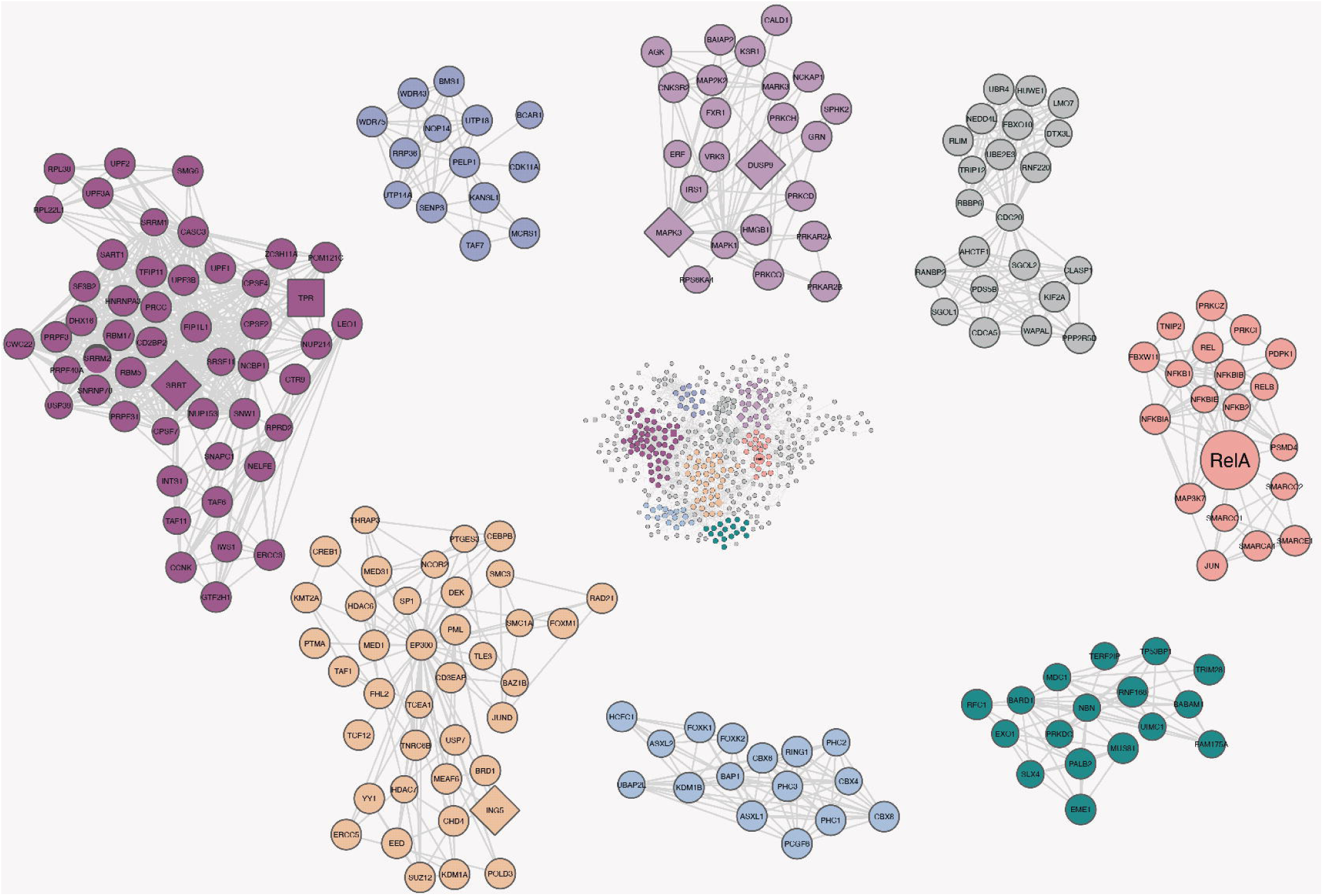

